# Subgenomic RNA Co-packaging Links Capsid Dynamics to Cell-to-Cell Movement in Brome Mosaic Virus

**DOI:** 10.64898/2026.02.02.703201

**Authors:** Antara Chakravarty, A. L. N. Rao

## Abstract

Subgenomic RNAs (sgRNAs) regulate gene expression in many positive-strand RNA viruses, yet their functional contributions beyond protein coding remain poorly understood. Here, we investigate how incorporation of subgenomic RNA4 (sgRNA4) into Brome mosaic virus (BMV) virions is associated with changes in virion structure, stability, and viral movement. Using an *in vivo* assembly system, we generated two distinct virion populations: particles containing RNA3 and sgRNA4 (B3+4^V^) and particles containing RNA3 alone (B3^V^). Although the two virion types were physically indistinguishable by electron microscopy, they exhibited strikingly different biochemical and biological properties. B3+4^V^ virions displayed increased protease sensitivity and structural flexibility, whereas B3^V^ virions were protease resistant and structurally rigid. MALDI–time of flight analysis revealed that trypsin digestion released similar peptide fragments from both virion types, while a persistent intact capsid protein peak in B3^V^ preparations indicated that only a small fraction of particles was accessible to proteolysis. In functional assays in *Nicotiana benthamiana*, B3+4^V^ virions were efficiently detected in adjacent tissues beyond the primary infiltration sites, whereas B3^V^ virions showed little to no cell-to-cell spread beyond the primary infiltration sites. Together, these results support a model in which sgRNA4 co-packaging is functionally linked to capsid dynamics that correlate with efficient viral movement. More broadly, our findings indicate that differences in packaged RNA composition can modulate virion physical properties and biological behavior, providing insight into how RNA viruses coordinate genome organization, assembly, and intercellular spread.

**Importance:** This study reveals that packaging of the small subgenomic RNA (sgRNA4) within BMV particles plays a crucial role beyond assembly—it is strongly associated with how the virus spreads between plant cells. Virions containing sgRNA4 are more flexible and dynamic, correlating with efficient movement through cell connections, while those lacking it are rigid and movement-deficient. These results uncover an unexpected link between RNA content and viral mobility, showing that genome composition can fine-tune both structural and biological properties of viruses—a principle that may extend to many other RNA viruses affecting plants and animals.

## Introduction

In many positive-sense single-stranded RNA viruses (such as BMV, Cucumber mosaic virus [CMV], or coronaviruses), the full-length genomic RNA serves a dual role as both the genetic material and the mRNA for replication-associated (nonstructural) proteins (1, 2). To expand their coding capacity and regulate gene expression, these viruses generate subgenomic RNAs (sgRNAs), which are typically produced either by internal initiation on negative-strand RNA templates or by discontinuous transcription (3–5). Both mechanisms are governed by cis-acting promoter elements embedded within the viral genome (3, 6). The modular arrangement of these promoters enables viruses to fine-tune the relative expression ratios of different proteins—a feature crucial for coordinated replication and assembly (3, 7).

Typically, colinear with the 3′ end of the genomic RNA, sgRNAs allow efficient translation of downstream open reading frames that cannot be accessed directly from the full-length genome due to ribosome scanning limitations (8). Functionally, sgRNAs uncouple genome replication from the synthesis of structural and movement proteins (3, 9). The full-length genomic RNA encodes replicase proteins required early in infection, whereas sgRNAs are synthesized later to drive high-level production of capsid and movement proteins (9, 10). This temporal separation ensures that genome amplification precedes virion assembly and intercellular spread (3, 9). In addition to structural proteins, some sgRNAs also encode accessory proteins that modulate host defense responses, viral movement, or pathogenesis (11).

In the *Bromoviridae*, particularly in BMV and CMV, sgRNAs play a central role in coordinating infection (12). BMV produces two sgRNAs: RNA4, which encodes the capsid protein, and RNA3-derived sgRNA3a, which encodes the movement protein (13, 14). The synthesis of these sgRNAs is tightly regulated by internal promoter elements on negative-strand replication intermediates, ensuring robust expression of structural and movement proteins only after replication complexes are established (13, 15).

Notably, BMV sgRNAs are not merely transcriptional outputs but are also frequently packaged into virions alongside full-length genomic RNAs (16, 17). Studies suggest that this encapsidation relies on conserved RNA motifs and tRNA-like structures at the 3′ ends, which serve multifunctional roles in translation, replication, and packaging (18, 19). The selective yet imperfect discrimination between genomic and subgenomic RNAs during encapsidation may introduce heterogeneity into viral populations, potentially conferring evolutionary flexibility (17, 20).

The packaging of sgRNAs is an intriguing and comparatively understudied phenomenon with implications for both viral biology and evolution (21). While most RNA viruses preferentially encapsidate genomic RNA to maximize infectivity, several viruses—including BMV, CMV, and Sindbis virus—also package sgRNAs, sometimes in significant proportions (16, 22). Encapsidation requires structural signals, including RNA stem-loop motifs, tRNA-like structures, and higher-order secondary/tertiary elements that interact with capsid proteins (23). Electrostatic interactions between positively charged capsid domains and sgRNA elements further stabilize particle formation (24). Multiple, non-mutually exclusive functions have been proposed for sgRNA-containing particles (21, 25). First, sgRNA-containing particles may act as defective or semi-infectious units that influence viral population dynamics, recombination, and genetic diversity (26). Second, packaging of sgRNAs could serve as an immune evasion strategy, producing noninfectious decoys that saturate host defenses (27). Third, sgRNAs may facilitate cross-packaging and heterologous recombination, driving viral evolution and host adaptation (26, 28). From a structural perspective, inclusion of sgRNAs may reflect the intrinsic flexibility of capsid–RNA recognition, in which multiple RNA substrates satisfy the geometric and charge requirements of assembly (24, 29).

This study investigates how the incorporation of the subgenomic RNA4 (sgRNA4) into BMV virions influences their structural dynamics and biological function. By comparing virions containing RNA3 and sgRNA4 (B3+4^V^) with those containing only RNA3 (B3^V^), we show that sgRNA4 packaging is not merely incidental but is functionally associated with differences in capsid dynamics and viral movement. Our results reveal that sgRNA4-containing particles exhibit enhanced capsid flexibility, increased susceptibility to proteolysis, and more efficient cell-to-cell movement, whereas particles lacking sgRNA4 are comparatively rigid, protease resistant, and movement-deficient. Together, these findings support a functional link between genome composition, capsid dynamics, and viral spread, uncovering a previously unappreciated role for sgRNA packaging in regulating plant RNA virus infectivity.

## Results

Throughout this study, the virion types B1^V^, B2^V^, B3^V^, and B3+4^V^ refer to individual virions that package genomic RNA1, RNA2, RNA3 only, and RNA3 together with RNA4, respectively. The wild-type (WT) represents a mixture of the three BMV virions—each containing RNA1, RNA2, and RNA3 plus RNA4—purified from infected *Nicotiana benthamiana* plants.

### Strategy for *in vivo* assembly of BMV virion types containing RNA3+4 and RNA3 only

Fig. 1A summarizes the characteristic features of transfer DNA (T-DNA) -based vectors engineered to express the three biologically active genomic RNAs (gRNAs) of BMV—pB1, pB2, and pB3—during transient expression in *N. benthamiana* plants. B3/P10 represents a variant of pB3 harboring a proline substitution for an arginine residue at position 10 within the N-terminal arginine-rich motif (N-ARM). The resulting capsid protein (CP) variant expressed from B3/P10 assembles into virions that package RNAs 1, 2, and 3 but are defective in packaging the subgenomic RNA4. Similarly, the T-DNA constructs illustrated in Fig. 1B were designed for the transient expression of the BMV replicase proteins 1a (p1a) and 2a (p2a).

**Fig. 1.**
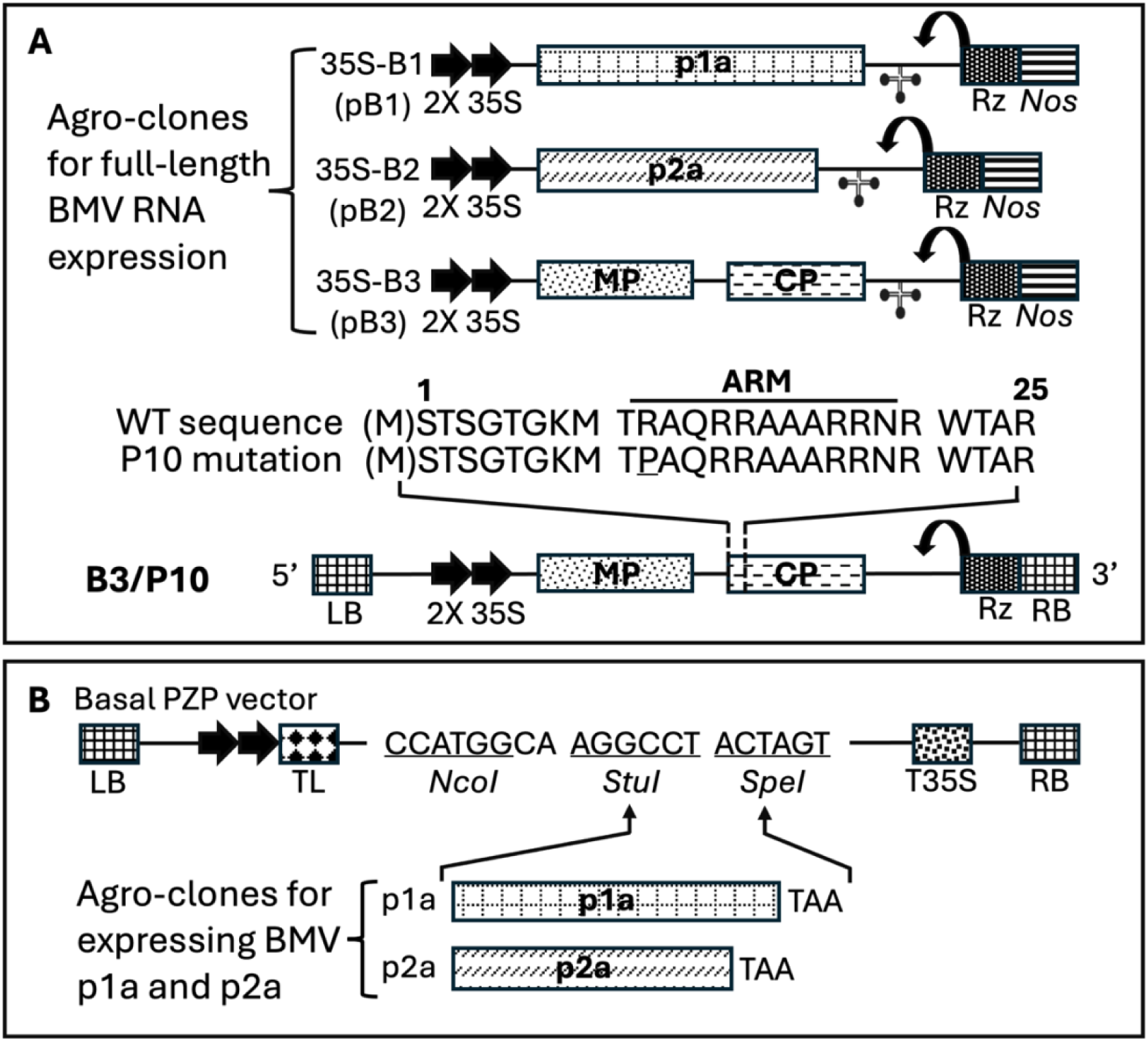
Characteristic features of agroplasmids used in this study. (A) Schematic representation of agroplasmids harboring Brome mosaic virus (BMV) genomic RNAs used for transient expression in plants. The 35S-B1 (pB1), 35S-B2 (pB2), and 35S-B3 (pB3) constructs contain full-length cDNA copies of BMV genomic RNAs 1 (B1), 2 (B2), and 3 (B3), respectively. B3/P10 denotes a mutant variant harboring a proline substitution for the 10th arginine residue in the N-terminal arginine-rich motif (N-ARM). The resulting capsid protein (CP) variant assembles into virions packaging RNAs 1, 2, and 3 but is defective in packaging sgRNA4. Single lines and boxes indicate noncoding and coding regions, respectively. Monocistronic B1 and B2 encode replicase proteins 1a (p1a) and 2a (p2a), respectively, while B3 contains open reading frames for the movement protein (MP) and CP. A cloverleaf-like structure at the 3′ end represents the conserved tRNA-like motif present in all three genomic RNAs. Double arrows at the 5′ ends indicate the duplicated 35S promoter, and a bent arrow marks the predicted self-cleavage site of the ribozyme. The location of the *nos* terminator is shown. (B) Agroconstructs for transient expression of p1a and p2a. Open reading frames (ORFs) of BMV p1a and p2a were cloned in-frame into binary vectors using *StuI* and *SpeI* restriction sites. Each binary vector contained, in order, the left border (LB) of T-DNA, a duplicated 35S promoter (35S×2), a tobacco etch virus (TEV) translational leader (TL), multiple cloning sites (MCS), a 35S terminator (T35S), and the right border (RB) of T-DNA (32, 57).

A schematic representation of the strategy used for *in vivo* assembly of virion types containing either RNA3 alone or RNA3 plus RNA4 is shown in Fig. 2. In formulating this strategy, three critical criteria were considered. First, genome packaging in bromoviruses is functionally coupled to replication—that is, only replication-derived progeny RNA is encapsidated (30, 31). Second, CP expressed in the absence of replication exhibits nonspecific RNA packaging activity (32). Third, packaging specificity is determined by an interaction between the BMV CP and the replicase protein 2a (33).

**Fig. 2.**
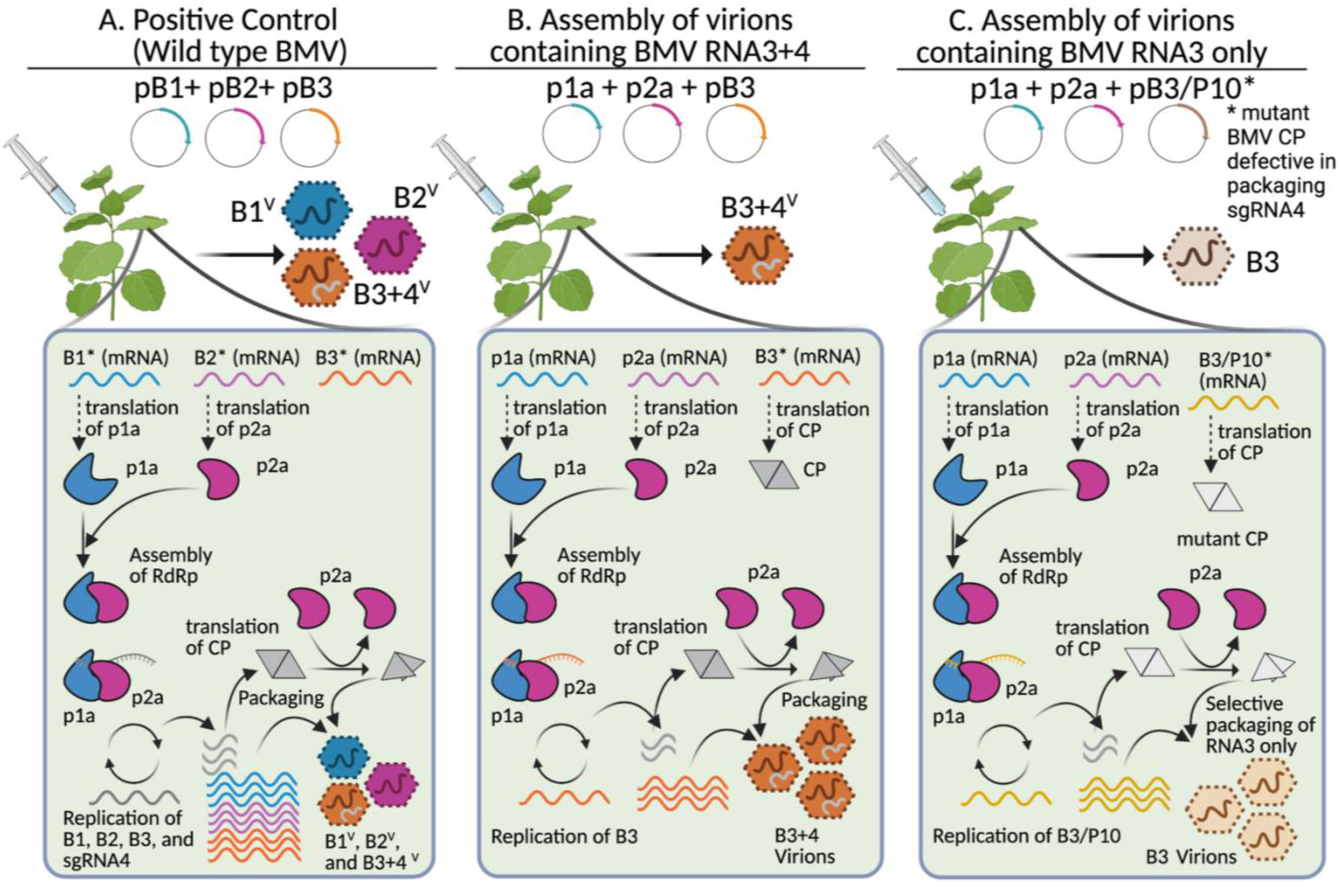
Schematic representation of the strategy used for autonomous in vivo assembly of Brome mosaic virus (BMV) virions differing in RNA composition. Infiltration of an inoculum containing a mixture of pB1, pB2, and pB3 results in WT BMV infection and the assembly of a mixture of all three virion types (B1^V^, B2^V^, and B3+4^V^). For the assembly of virions packaging RNA3 and sgRNA4 (B3+4^V^), an inoculum containing agrotransformants of p1a, p2a, and pB3 was infiltrated into *Nicotiana benthamiana* leaves. Transcription of p1a and p2a produces mRNAs competent for translation but not replication due to the absence of 5′ and 3′ noncoding regions. Translation yields replicase proteins 1a and 2a, which assemble into a functional replicase complex that directs replication of B3 RNA and synthesis of sgRNA4 for capsid protein (CP) production. The CP subunits translated from sgRNA4 subsequently package both B3 RNA and sgRNA4 into virions, forming B3+4^V^ particles. In contrast, for the assembly of virions packaging only RNA3 (B3^V^), an inoculum containing p1a, p2a, and pB3/P10 was used (39). Replication of pB3/P10 results in sgRNA4 synthesis and CP production, but the P10 mutation disrupts the interaction between the N-terminal arginine-rich motif (N-ARM) and sgRNA4, thereby preventing sgRNA4 encapsidation. Consequently, B3^V^ virions package only RNA3 and are defective in sgRNA4 packaging. Created in BioRender. Chakravarty, A. (2026) https://BioRender.com/89hgduo.

Taking these considerations into account, the strategy outlined in Fig. 2 was designed to direct the assembly of desired virion types by agroinfiltration with the following inocula:

(i) *Control assembly.* Plants infiltrated with a mixture of agrotransformants expressing all three full-length RNAs (pB1, pB2, and pB3) (Fig. 2A). (ii) *Assembly of RNA3+4 virions (B3+4^V^).* An inoculum comprising agrotransformants p1a, p2a, and pB3 (Fig. 2B). Upon infiltration, p1a and p2a transcripts serve as mRNAs for the synthesis of 1a and 2a proteins, respectively. Although these mRNAs lack the 5′ and 3′ noncoding regions required for replication (Fig. 1B), transiently expressed 1a and 2a associate to form a functional replicase that replicates RNA3 and produces sgRNA4 for CP expression, resulting in the assembly of B3+4^V^ particles. (iii) *Assembly of RNA3-only virions (B3^V^).* An inoculum consisting of p1a, p2a, and pB3/P10 (Fig. 2C). Replication of pB3/P10 leads to the synthesis of sgRNA4 for CP production, but the P10 mutation disrupts the interaction between N-ARM and sgRNA4, thereby preventing sgRNA4 encapsidation. Consequently, the resulting virions package only RNA3.

### Characteristic features of B3+4^V^ and B3^V^

Purified virions obtained from *N. benthamiana* leaves infiltrated with either the WT control inoculum (pB1 + pB2 + pB3) (Fig. 2A) or with p1a + p2a + pB3 (B3+4^V^; Fig. 2B) or p1a + p2a + pB3/P10 (B3^V^; Fig. 2C) were examined by negative-stain electron microscopy (EM). As shown in Fig. 3 (A-C), virions derived from both B3+4^V^ and B3^V^ infiltrations were morphologically indistinguishable from those of the WT control (a mixture of B1^V^, B2^V^, and B3+4^V^), exhibiting uniform, spherical particles with an average diameter of approximately 28 nm. Northern blot analysis confirmed that the WT, B3+4^V^, and B3^V^ virions encapsidated the expected RNA progeny (data not shown).

**Fig. 3.**
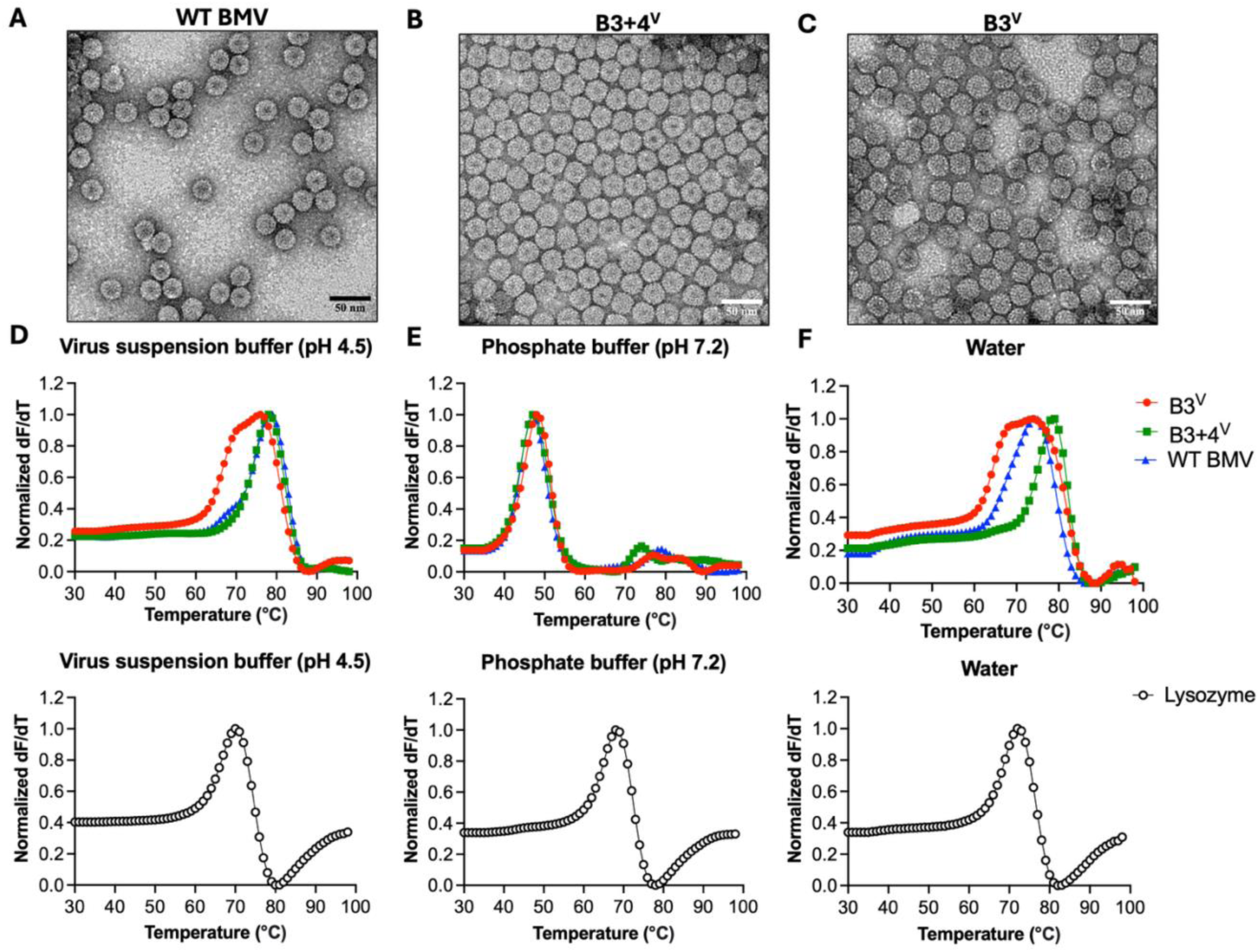
Morphology and thermal stability of WT BMV, B3+4^V^, and B3^V^ virions. (A-C) Negative-stain EM images showing the virion preparations for the indicated samples. Scale bar: 50 nm. Representative results are shown from *n = 2* independent experiments. (D-F) Stability analysis of *Brome mosaic virus* (BMV) B3+4^V^, B3^V^, and wild-type (WT) virions by differential scanning fluorimetry (DSF). The first derivative of fluorescence intensity is plotted as a function of temperature for each virion type, normalized so that the maximum derivative equals 1 on the y axis. Thermal denaturation profiles of B3+4^V^, B3^V^, and WT virions were recorded under the indicated buffer and pH conditions, with lysozyme included as a positive control. The virions of B3+4^V^, B3^V^, and WT BMV exhibited nearly identical temperature-dependent melting transitions (∼70°C) in sterile water (pH 7.1) and virus suspension buffer (pH 4.5), although B3^V^ displayed a broader melting peak, initiating denaturation slightly earlier than B3+4^V^ and WT BMV. In 100 mM phosphate buffer (pH 7.2), all three virion types showed reduced thermal stability, melting primarily at ∼50°C. Lysozyme exhibited a consistent melting profile (∼60°C) under all three conditions, confirming assay uniformity. Representative results are shown from *n = 3* independent experiments.

### Stability of B3+4^V^ and B3^V^ virions

To assess the relative thermal stability of B3+4^V^ and B3^V^ virions, differential scanning fluorimetry (DSF) was performed as described in Materials and Methods. The hydrophobic dye SYPRO Orange binds to hydrophobic regions of CP exposed during thermal denaturation, and the resulting increase in fluorescence intensity reflects progressive CP unfolding (34). DSF profiles depicting temperature-dependent melting of B3+4^V^, B3^V^, and control samples (WT BMV assembled in *N. benthamiana* and lysozyme) under three buffer and pH conditions are summarized in Fig. 3 (D-F). In virus suspension buffer (pH 4.5), all virion types exhibited comparable thermal stability, with melting transitions occurring near 70°C (Fig. 3D, top panel). B3^V^ displayed a broader melting peak, initiating denaturation slightly earlier than B3+4^V^ and WT BMV. In phosphate buffer (pH 7.2), two thermally unstable populations were observed at approximately 50°C and 70°C, with all virion types showing a major melting peak at ∼50°C (Fig. 3E, top panel). Similarly, in water (pH 7.1), B3+4^V^, B3^V^, and WT BMV exhibited comparable melting behavior (Fig. 3F, top panel). Under these conditions, B3+4^V^ melted near 75°C, slightly higher than the ∼70°C melting point of B3^V^ and WT BMV, consistent with previous observations (35, 36). Lysozyme showed no significant variation across buffer conditions (Fig. 3D-F, bottom panel).

### Comparative structural dynamics of B3+4^V^ and B3^V^ virions revealed by proteolysis and mass spectrometry

Limited trypsin proteolysis followed by Western blotting, transmission electron microscopy (TEM), and mass spectrometry was used to evaluate virion structural dynamics. Each virion preparation was divided into two aliquots after digestion: one analyzed by Western blotting with anti-CP antibody, the other by TEM to assess virion integrity.

Undigested virions of B3+4^V^, B3^V^, and WT BMV migrated as single, intact CP bands (∼20 kDa), with monomeric and dimeric forms detected in all samples (Fig. 4A, lanes 1). Trypsin digestion produced strikingly distinct profiles. B3+4^V^ virions were highly susceptible, showing extensive degradation within 10 min (Fig. 4A, lanes 2–5). In contrast, B3^V^ virions displayed strong protease resistance, retaining intact CP throughout digestion (Fig. 4A). WT BMV showed a digestion pattern similar to that of B3+4^V^, consistent with the predominance of B3+4^V^ in WT preparations (35). Densitometric analysis indicated that ∼68% of CP in B3+4^V^ and WT was digested by trypsin, while ∼87% of CP in B3^V^ remained intact after 3 h.

**Fig. 4.**
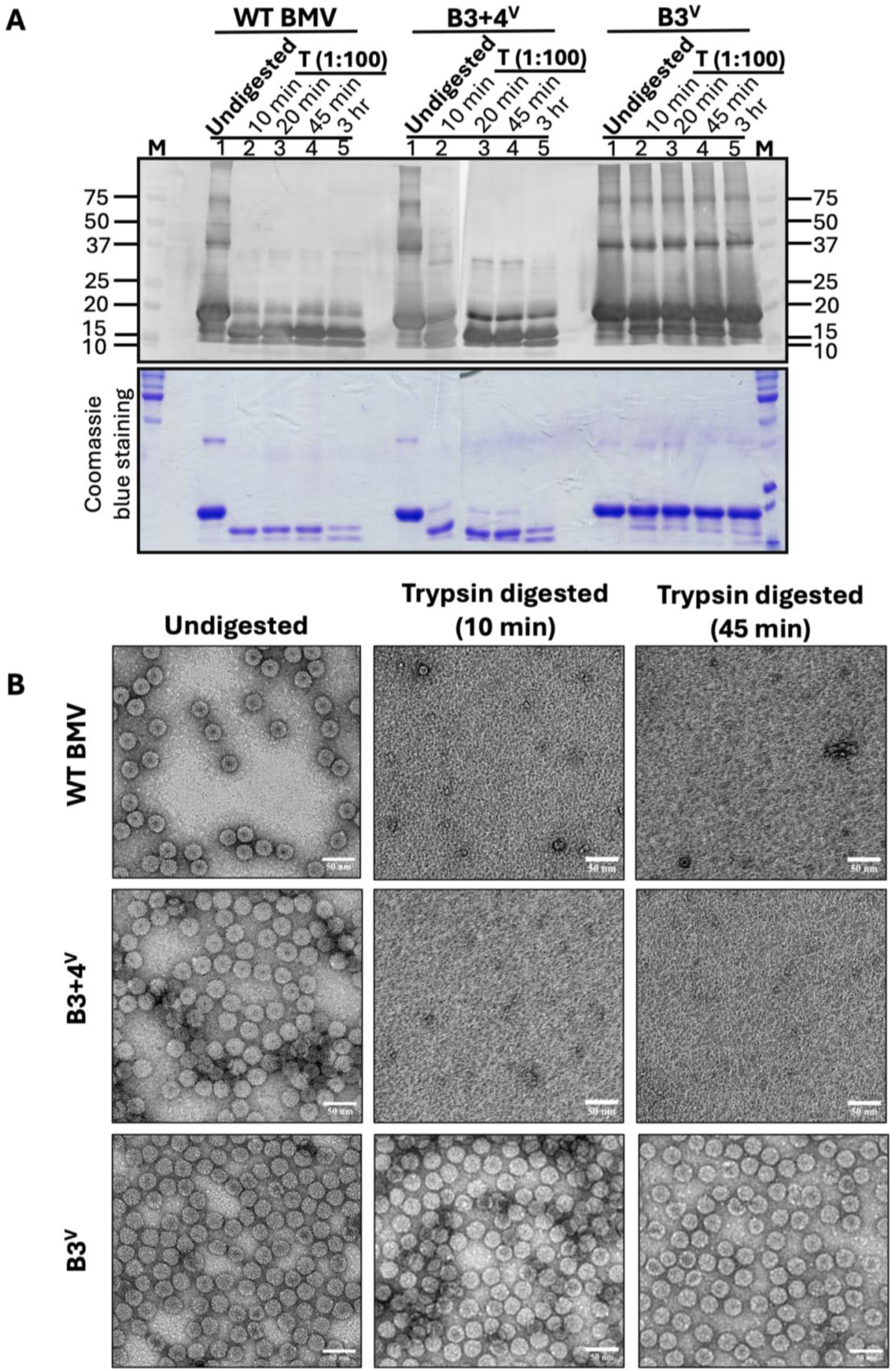
Differential susceptibility of B3+4V and B3V virions to limited trypsin proteolysis. (A) Western blot analysis of B3+4^V^, B3^V^, and WT BMV virions following limited trypsin digestion. Each virion preparation was either left undigested or treated with trypsin for 10, 20, or 45 min or for 3 h prior to electrophoresis. (B) Negative-stain EM images showing the structural integrity of undigested and trypsin-digested virion preparations for the indicated samples. Scale bar: 50 nm. Representative results are shown from *n = 2* independent experiments.

TEM confirmed these results (Fig. 4B). Undigested virions were uniformly spherical (∼28 nm). After trypsin treatment, B3+4^V^ and WT virions appeared disrupted or collapsed, while B3^V^ virions retained icosahedral morphology even after extended incubation, consistent with a more protease-resistant and structurally constrained capsid state.

MALDI–time of flight (TOF) mass spectrometry further characterized proteolytic fragments (Fig. 5-7). Undigested preparations yielded a single peak (∼20 kDa), confirming homogeneity and purity (Fig. 5A). Trypsin-digested B3+4^V^ samples exhibited multiple lower-mass fragments that increased with digestion time (Fig. 5B), reflecting surface-exposed cleavage sites. By contrast, B3^V^ displayed a similar number of cleavage products indicating very few trypsin-susceptible virions (Fig. 6A), and the intact 20-kDa peak persisted after 3 h, indicating reduced accessibility and a more compact capsid (Fig. 6B).

**Fig. 5.**
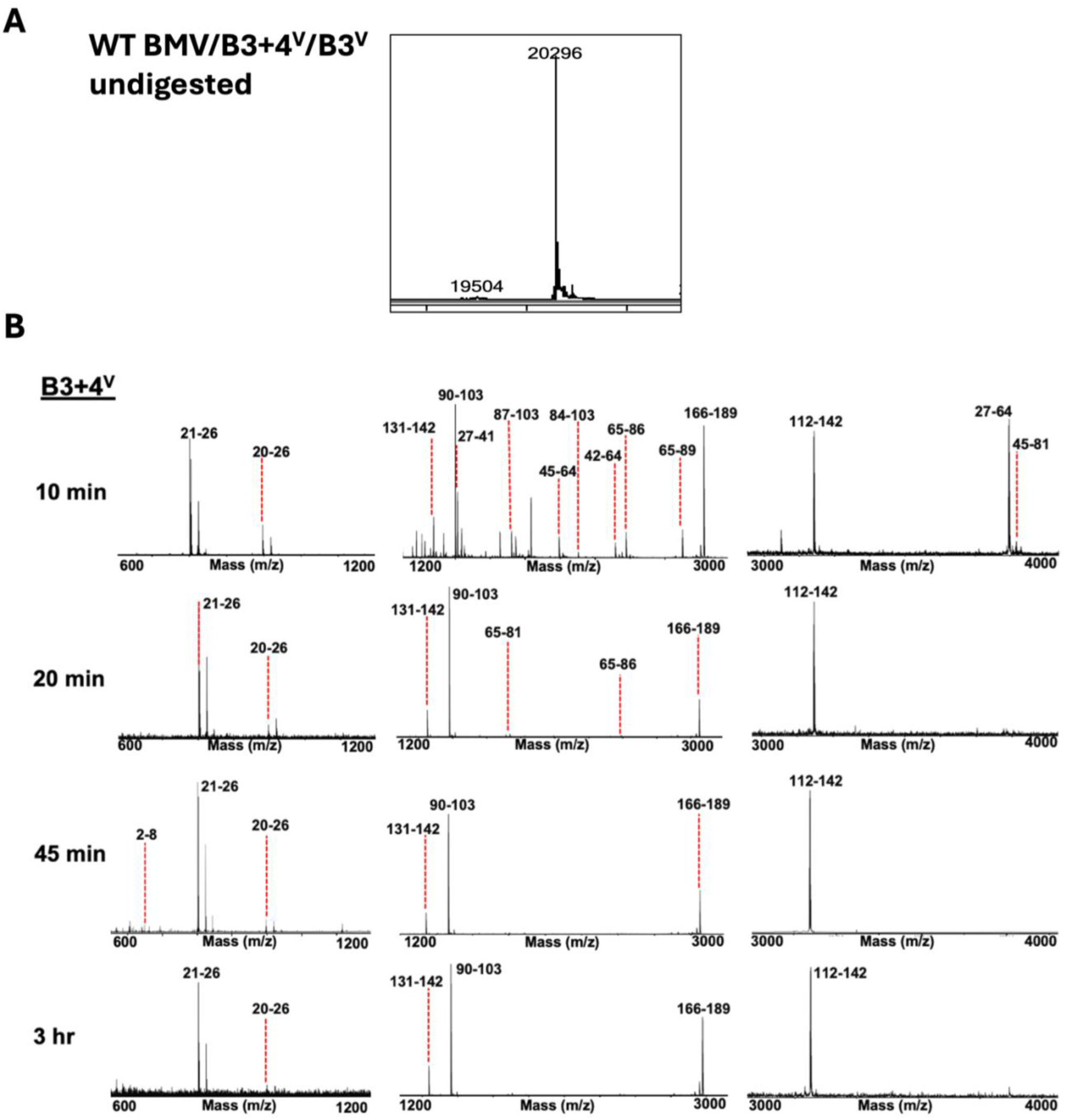
Matrix-assisted laser desorption ionization–time of flight (MALDI-TOF) mass spectrometric analysis of undigested and trypsin-digested Brome mosaic virus (BMV) virions. (A) Mass spectra of undigested B3^V^, B3+4^V^, and wild-type (WT) BMV virions, each displaying a single predominant peak at ∼20,296 Da, corresponding to the intact capsid protein. (B) MALDI–TOF mass spectrometric analysis of capsid protein fragments released from sgRNA4-containing virions following limited proteolysis. Peaks are labeled with the corresponding capsid protein fragments and their predicted amino acid residue positions. Representative results are shown from *n = 2* independent experiments.

**Fig. 6.**
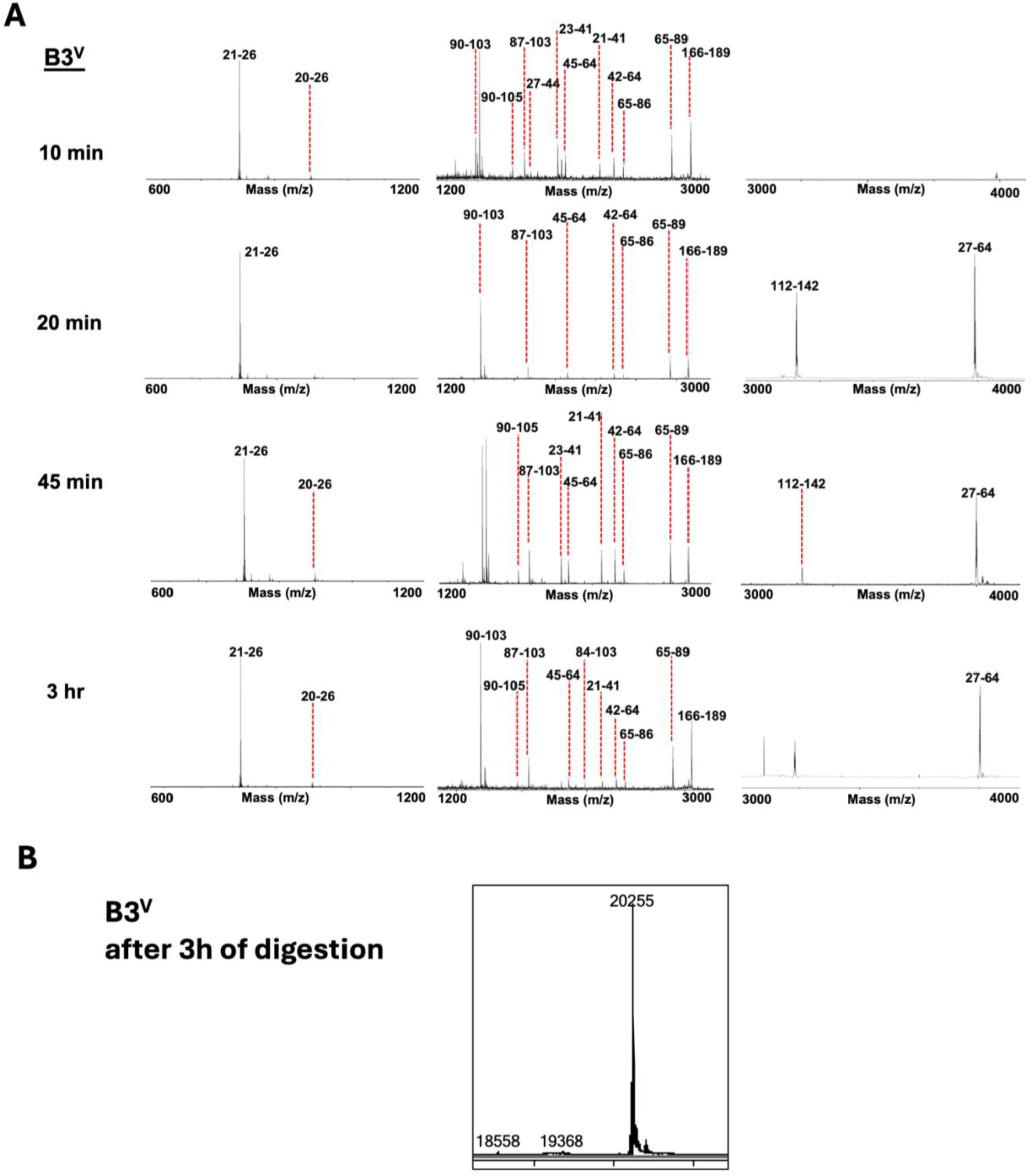
Persistence of intact capsid protein in RNA3-only virions following extended trypsin digestion. (A) MALDI-TOF analysis of peptides released from B3^V^ virions at the indicated time points following trypsin digestion. Peaks are labeled with the corresponding capsid protein fragments and their predicted amino acid residue positions. (B) LC-MS analysis of B3^V^ after 3 h of trypsin digestion revealed a ∼20kDa peak. Representative results are shown from *n = 2* independent experiments.

**Fig. 7.**
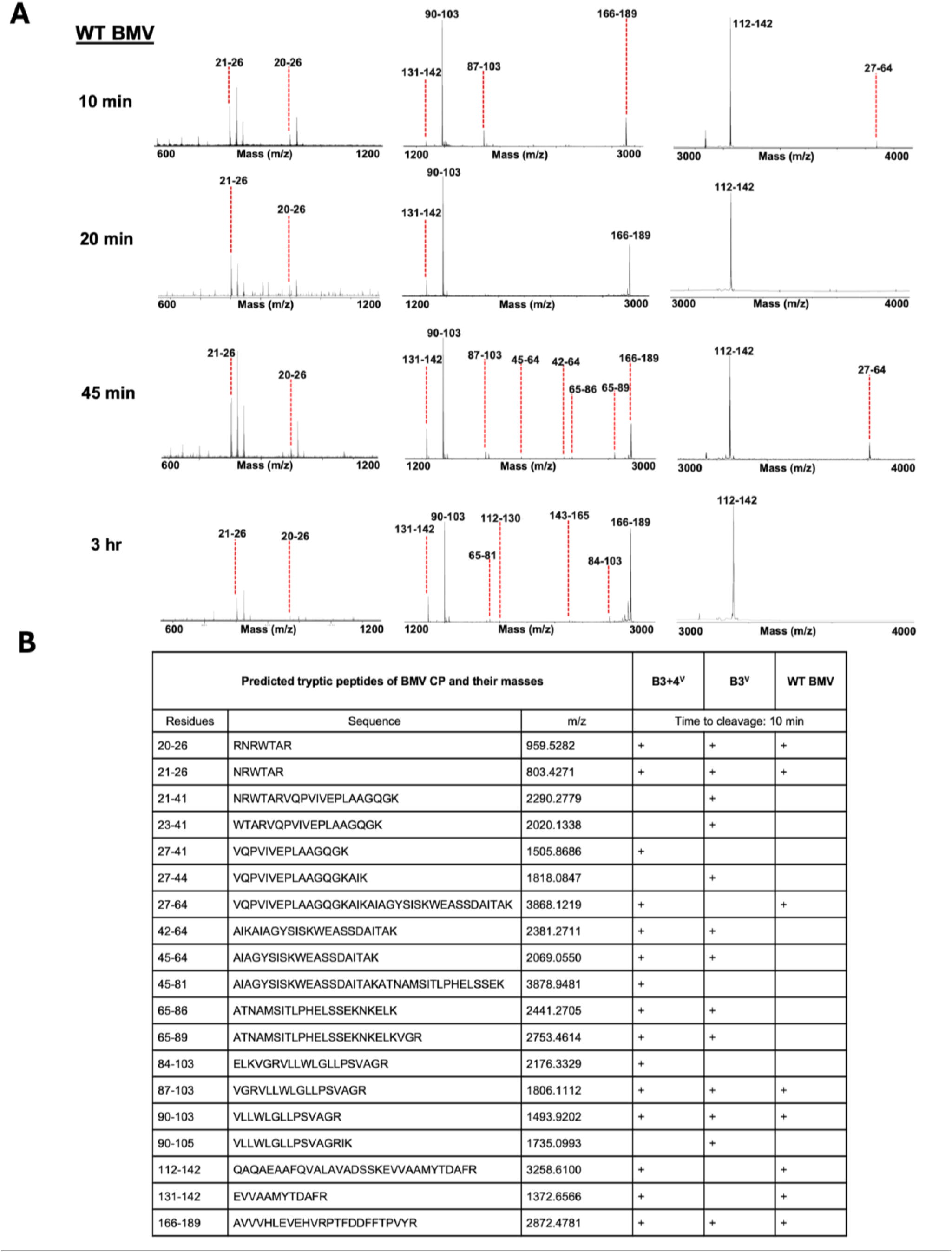
Proteolytic profile of wild-type BMV virions following limited trypsin digestion. (A) MALDI-TOF analysis of peptides released from wild-type (WT) Brome mosaic virus (BMV) virions at the indicated time points following trypsin digestion. Peaks are labeled with the corresponding capsid protein fragments and their predicted amino acid residue positions. (B) Table showing comparison of tryptic peptides derived from B3+4^V^, B3^V^, and WT BMV after 10 min of trypsin digestion. Representative results are shown from *n = 2* independent experiments.

Collectively, Western blot, TEM, and MALDI-TOF data revealed that B3+4^V^ and WT BMV (Fig. 7A), possess dynamic, protease-sensitive capsids, whereas B3^V^ forms rigid, protease-resistant structures. The presence of sgRNA4 thus contributes to enhanced capsid flexibility, underscoring the role of RNA composition in modulating virion stability and dynamics.

### Influence of sgRNA4 packaging on virus spread

In *Chenopodium quinoa*, chlorotic and necrotic lesion spread complicates visualization of infection boundaries (37), whereas *N. benthamiana* infected with BMV exhibits no visible symptoms, making it ideal for studying local movement (38). Distinct movement patterns were observed between WT BMV and pB1 + pB2 + pB3/P10 (B3^V^) infections. In plants infiltrated with pB1 + pB2 + pB3 (WT), CP expression was detected in both source and sink regions by 4 days post-infiltration (dpi), persisting through 10 dpi (Fig. 8A, B). In contrast, pB1 + pB2 + pB3/P10 infiltration yielded reduced CP accumulation, detectable in sink regions only up to 4 dpi, and later restricted to the source region (Fig. 8B). Notably, reduced CP accumulation in source tissues in pB1 + pB2 + pB3/P10 infections relative to WT at early time points, suggested that the absence of sgRNA4 affects the establishment or amplification of CP expression in planta. Because CP accumulation in intact leaf tissue integrates replication, subgenomic RNA production, and cell-to-cell spread, differences in CP abundance are not expected to strictly mirror those observed in single-cell expression systems (39). To determine whether impaired movement of virions lacking sgRNA4 reflected defects in genome dissemination, we performed RT–PCR analysis on total RNA isolated from source and sink tissues at 4 dpi. In plants infected with WT BMV (pB1 + pB2 + pB3), robust accumulation of BMV RNA1 and RNA2 was detected in source regions, accompanied by strong signals for MP and CP transcripts (Fig. 8C). Similarly, virions containing RNA3 and sgRNA4 (B3+4^V^) supported efficient detection of genomic RNAs and sgRNA-derived transcripts in sink tissues, consistent with productive cell-to-cell movement. In contrast, infections with virions lacking sgRNA4 (B3^V^) showed only weak MP RNA accumulation in source tissue and failed to yield detectable MP or CP signals in sink regions, despite the presence of RNA1 and RNA2. These observations are consistent with the possibility that the rigid B3/P10 capsids limit processes required for productive infection beyond the initially infected cells. The detection of RNA1 and RNA2 in sink tissues under both conditions indicates that viral replication-associated RNA dissemination can occur independently of sgRNA4, whereas productive accumulation of movement- and capsid-associated transcripts in distal cells requires sgRNA4 copackaging. Together with the protein-level defects observed by immunoblotting, these findings demonstrate that sgRNA4 packaging is functionally coupled to cell-to-cell movement by coordinating capsid dynamics with localized expression of movement-associated viral functions.

**Fig. 8.**
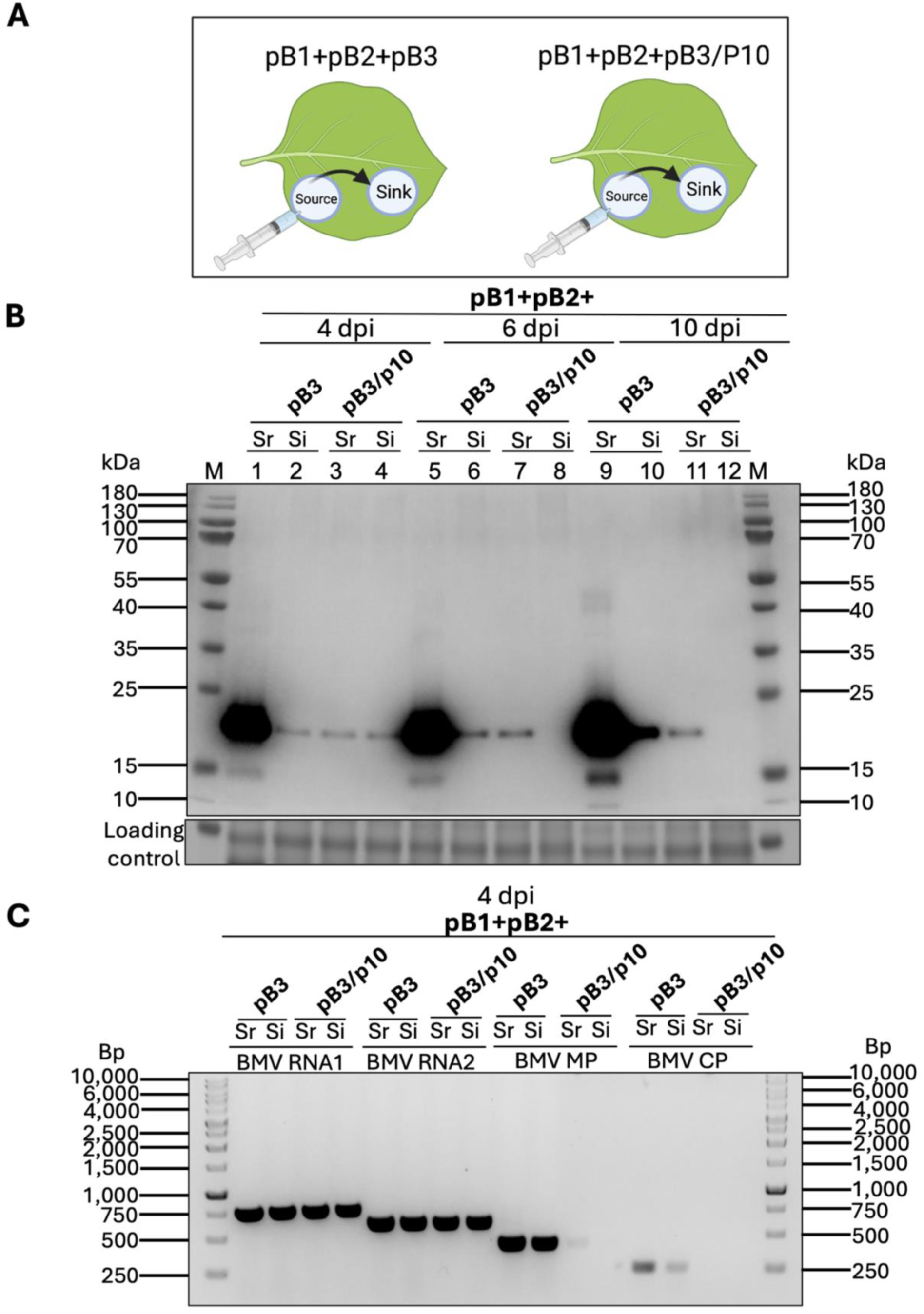
Accumulation and distribution of viral gene products in source and sink tissues following infection with sgRNA4-containing or sgRNA4-deficient virions. (A) Schematic representation of the experimental setup showing the relative positions of the agroinfiltrated (source) and non-infiltrated (sink) leaf discs used for analysis. Created in BioRender. Chakravarty, A. (2026) https://BioRender.com/4dx1p4g. (B) Western blot analysis (top panel) shows the relative accumulation of BMV capsid protein (CP) in samples loaded in lanes 1–12 as indicated. SDS-PAGE analysis of total protein extracts stained with Coomassie Brilliant Blue R-250 (bottom panel) illustrates the relative accumulation of viral proteins in source and sink regions at 4, 6, and 10 days post-infiltration (dpi) with the indicated inocula. Each lane contained 30 µg of total protein, as determined by the Bradford assay. (C) RT–PCR analysis of viral RNAs isolated from source (Sr) and sink (Si) leaf tissues at 4 days post-infiltration (dpi) following inoculation with pB1 + pB2 in combination with pB3 or pB3/P10. Total RNA was extracted from the indicated tissues and analyzed using BMV-specific primers to detect genomic RNAs (RNA1 and RNA2) as well as subgenomic RNA–encoded transcripts corresponding to the movement protein (MP) and capsid protein (CP). PCR products were resolved on agarose gels, with molecular size markers (bp) shown. Robust amplification of RNA1 and RNA2 was detected in both source and sink tissues under both conditions, whereas MP and CP transcripts were readily detected in sink tissues only when sgRNA4-containing virions were present (pB3), consistent with efficient dissemination to adjacent cells. In contrast, infections with virions lacking sgRNA4 (pB3/P10) showed reduced MP signal in source tissue and no detectable MP or CP transcripts in sink regions, consistent with impaired viral spread. These assays report tissue-level distribution of viral gene products and do not directly visualize virion transit between individual cells. Representative results are shown from *n = 2* independent experiments.

Since sgRNA4 encodes CP, its co-packaging with RNA3 is associated with enhanced capsid flexibility and a property consistent with efficient virion trafficking through plasmodesmata. The rigid, protease-resistant capsid of B3^V^ likely limits this process, reducing spread beyond initially infected cells. These findings support a model in which sgRNA4 packaging modulates capsid plasticity—balancing structural stability with dynamic movement, essential for efficient intercellular transport and systemic infection. Together, these findings reveal consistent biochemical, structural, and biological differences between sgRNA4-containing and sgRNA4-deficient virions, which are further interpreted in the Discussion.

## Discussion

In this study, we investigated how the composition of encapsidated RNA influences the structural properties and biological behavior of BMV virions. By using an *in vivo* assembly strategy to generate virions containing RNA3 together with subgenomic RNA4 (B3+4^V^) or RNA3 alone (B3^V^), we were able to directly compare particles that differ primarily in their packaged RNA content (Fig. 1, 2). Our results show that sgRNA4 copackaging is associated with pronounced differences in capsid dynamics, protease sensitivity, and the capacity for cell-to-cell movement, supporting a functional relationship between virion RNA composition and viral cell-to-cell movement.

### Influence of sgRNA4 Packaging on Virion Structural Dynamics

Despite appearing morphologically similar by electron microscopy, B3+4^V^ and B3^V^ displayed distinct physicochemical properties. Differential scanning fluorimetry revealed broadly comparable thermal stability under multiple buffer conditions (Fig. 3), while limited proteolysis and mass spectrometry uncovered striking differences in capsid dynamics. B3+4^V^ and WT virions were readily susceptible to trypsin digestion (Fig. 4, 5, 7), whereas B3^V^ particles remained largely intact, retaining a full-length 20-kDa CP peak even after extended proteolysis (Fig. 4, 6). These findings are consistent with a more compact and protease-resistant capsid architecture in B3^V^ particles and a comparatively flexible or dynamic capsid conformation in B3+4^V^ virions.

The observation that similar tryptic peptides were released from both virion types suggests that the underlying capsid structures are not fundamentally altered but instead differ in the extent to which cleavage sites are transiently exposed. Such behavior is characteristic of metastable viral capsids, in which localized conformational dynamics permit controlled accessibility without global disassembly. Similar RNA-dependent modulation of capsid dynamics has been described for other icosahedral viruses, where encapsidated RNA contributes to capsid expansion, uncoating, or mechanical flexibility required during infection (40–42). In this context, sgRNA4 packaging may influence the conformational landscape of the BMV capsid, favoring a dynamic state compatible with downstream biological functions.

### Functional coupling of virion dynamics to viral movement

The structural differences observed between B3+4^V^ and B3^V^ virions correlated with marked differences in their ability to move between cells. In *Nicotiana benthamiana*, infections involving wild-type or B3+4^V^ virions resulted in sustained accumulation of capsid protein in both source and sink tissues, whereas infections with sgRNA4-deficient virions showed reduced and transient capsid protein accumulation that remained largely confined to source regions. Consistent with these protein-level observations, RT–PCR analysis revealed efficient accumulation of movement- and capsid-associated transcripts in sink tissues only when sgRNA4-containing virions were present.

The capsid protein of plant viruses is increasingly recognized as a multifunctional component that bridges replication, assembly, and intercellular movement (31, 43). Beyond its classical role in genome encapsidation, the CP actively participates in the formation of transport-competent complexes and regulates virion adaptability during infection (44–46). Specific domains, particularly the N-terminal arginine-rich motifs (N-ARMs), mediate RNA binding and membrane targeting, while conformational flexibility enables virions to transiently expand or partially disassemble as they traffic through plasmodesmata (47–50). This structural plasticity is essential for balancing genome protection with the need for dynamic interactions with host and viral factors (51, 52). In plant RNA viruses such as BMV, CMV, and *Tobacco mosaic virus* (TMV), efficient movement relies on a finely tuned equilibrium between capsid rigidity and flexibility—features that permit passage through plasmodesmata without premature disassembly (53–55). Thus, virion formation is not merely a terminal event of infection but an integral determinant of viral dissemination and host adaptation. The capsid protein also functions as a regulatory element that coordinates replication-derived packaging with movement processes (31). Mutations that alter CP flexibility or disrupt RNA–protein and CP–MP interactions frequently result in defective systemic spread (49, 53, 56). Conversely, the presence of sgRNA4 within B3+4^V^ virions appears to preserve the metastable state necessary for controlled uncoating and interaction with movement machinery (Fig. 9). These findings reinforce the concept that virion assembly, structural dynamics, and movement are tightly interdependent events within the viral life cycle.

**Fig. 9.**
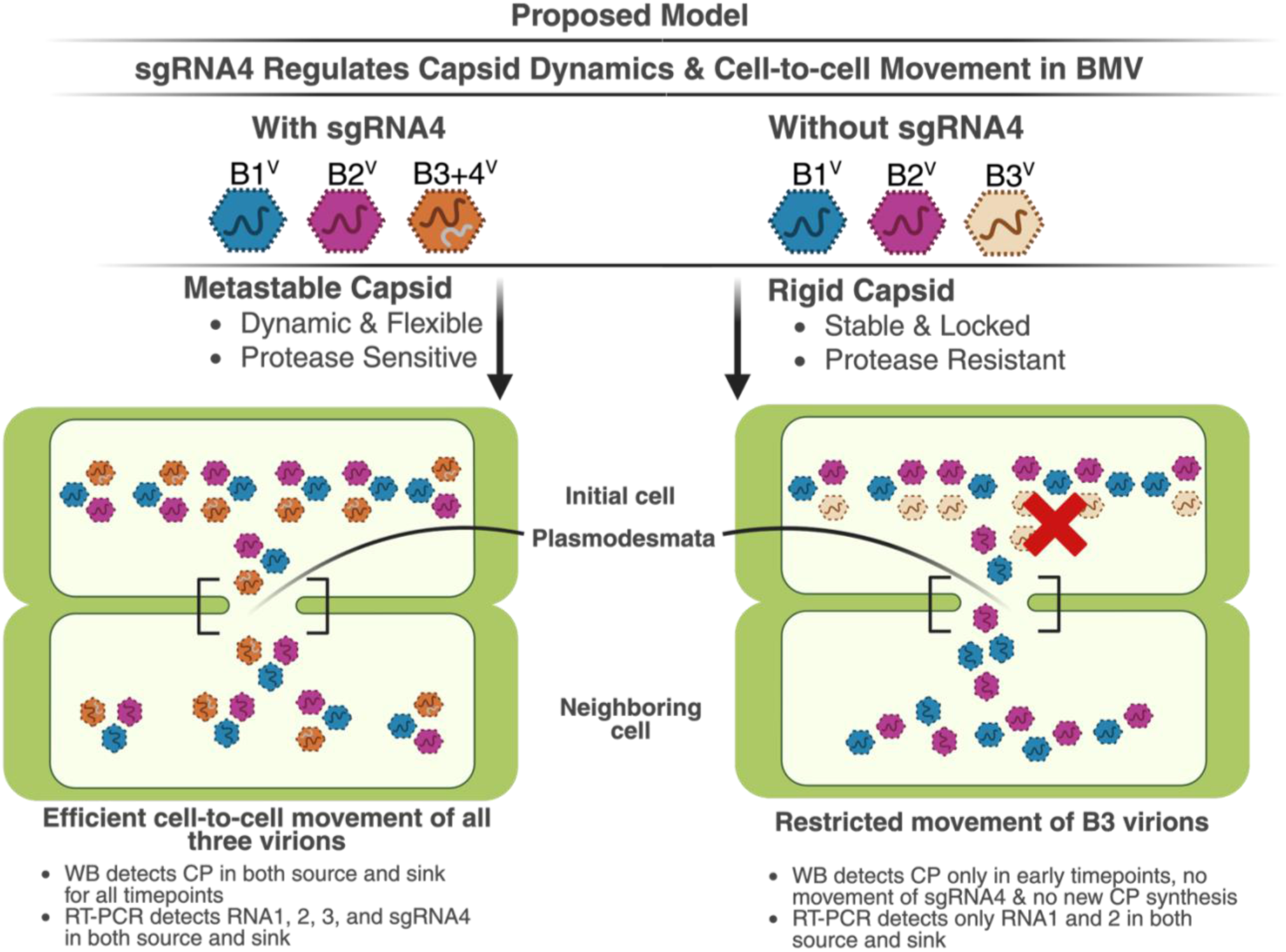
Proposed model linking sgRNA4 co-packaging to capsid dynamics and cell-to-cell spread in Brome mosaic virus. Schematic representation of how copackaging of subgenomic RNA4 (sgRNA4) with RNA3 influences virion capsid properties and intercellular movement. In the presence of sgRNA4 (left), BMV virions containing RNA1 (B1^V^), RNA2 (B2^V^), and RNA3 copackaged with sgRNA4 (B3+4^V^) adopt a metastable capsid state characterized by structural flexibility and protease sensitivity. This metastable configuration enables efficient trafficking of all three virion types through plasmodesmata from initially infected source cells to adjacent sink cells, resulting in robust accumulation of capsid protein (CP) and viral RNAs in both tissues. In the absence of sgRNA4 packaging (right), virions containing RNA1 and RNA2 retain the capacity for cell-to-cell movement, whereas RNA3-only virions (B3^V^) form rigid, protease-resistant capsids that are defective for movement and/or genome uncoating. Consequently, CP accumulation and viral RNA detection in sink tissues are restricted, reflecting impaired dissemination of RNA3-derived functions. Together, the model illustrates how sgRNA4 copackaging modulates capsid dynamics to functionally couple virion assembly with intercellular movement during BMV infection. This model summarizes functional associations supported by the present data and does not imply a direct causal mechanism. Created in BioRender. Chakravarty, A. (2026) https://BioRender.com/gtogrc2.

Unlike T7-driven expression in barley protoplasts that study intracellular protein expression in isolated cells (39), in intact plant tissue, capsid protein accumulation reflects the integrated outcomes of replication, subgenomic RNA production, intercellular movement, and the establishment of infection in neighboring cells. Thus, reduced capsid protein accumulation in sink tissues in the absence of sgRNA4 is most consistent with a defect in productive cell-to-cell spread rather than a primary defect in intracellular replication. Notably, viral RNAs encoding replication functions were detected in sink tissues under both conditions, indicating that dissemination of replication-associated RNA can occur independently of sgRNA4 packaging, whereas robust accumulation of movement- and capsid-associated gene products correlates with the presence of sgRNA4-containing virions. Because CP accumulation in WT source tissue approached saturation under the exposure conditions used, immunoblot signal intensity is not strictly quantitative across conditions. Importantly, however, the absence of detectable CP in sink tissues at later time points in pB1 + pB2 + pB3/P10 infections was reproducible across experiments, indicating a genuine defect in CP accumulation and spread rather than an artifact of antibody saturation.

### Proposed mechanistic model and implications for virus biology

Although sgRNAs are classically viewed as transcriptional intermediates that direct the synthesis of structural and movement proteins, increasing evidence indicates that they play additional roles in viral assembly and infection. Our findings support a model in which sgRNA4 co-packaging is functionally linked to virion properties that influence biological outcomes, rather than serving solely as a passive byproduct of replication. Importantly, while the present data do not formally establish a direct causal mechanism linking sgRNA4 packaging to virion mobility, the convergence of biochemical, structural, and biological phenotypes strongly supports a model in which sgRNA4 contributes to a metastable capsid state that correlates with efficient cell-to-cell movement. Definitive separation of RNA packaging effects from downstream translational or replication-associated consequences will require future experimental approaches beyond the scope of the current study.

A schematic model summarizing this relationship is presented in Fig. 9. In this model, co -packaging of sgRNA4 with RNA3 favors a capsid conformation that balances stability with dynamic adaptability, enabling virions to traffic between cells and support productive infection. In contrast, virions lacking sgRNA4 adopt a more rigid conformation that correlates with impaired movement and reduced accumulation of movement- and capsid-associated gene products in distal tissues. Future studies that uncouple sgRNA packaging from downstream translational effects will be required to further refine this model and define the molecular mechanisms involved.

Together, these results highlight sgRNAs as multifunctional regulatory elements whose roles extend beyond protein coding to include modulation of virion properties and infection dynamics. By influencing capsid behavior, packaged sgRNAs may contribute to the coordination of genome organization, assembly, and movement that underpins efficient viral infection. This work underscores the importance of RNA composition as a determinant of virion function and suggests that RNA-dependent modulation of capsid dynamics may represent a general strategy employed by RNA viruses to optimize infectivity and spread.

## Materials and methods

### Agroplasmids used in this study

The construction and characteristic features of agroplasmids pB1, pB2, and pB3 (Fig. 1A), engineered to express biologically active, full-length genomic RNAs of BMV following agroinfiltration into plants, have been described previously (32). The plasmid pB3/P10 represents a B3 variant clone containing a proline substitution for an arginine residue at position 10 within the N-terminal arginine-rich motif (N-ARM), resulting in the production of virions defective in sgRNA4 packaging (39). Similarly, agroplasmids p1a and p2a (Fig. 1B) were constructed to transiently express the BMV replicase proteins 1a and 2a, respectively, as described previously (32, 57).

### Agroinfiltration, virion purification, EM, and Western blot analysis

All agroplasmids were introduced into *Agrobacterium tumefaciens* strain GV3101. The transformed cultures were infiltrated at identical optical densities into the abaxial surface of wild-type *Nicotiana benthamiana* leaves as described previously (58), and equal amounts of total protein or RNA were used for all downstream analyses. WT, B3+4^V^, and B3^V^ virions were purified from 4-day-post-infiltration (dpi) leaves using the same procedure employed for WT BMV, followed by sucrose density gradient centrifugation (59). For negative-stain EM, virions were imaged using a Tecnai 12 TEM operated at 120 keV, and images were recorded digitally. Western blot analyses of undigested and trypsin-digested virion samples were performed using anti-CP antibodies as described previously (32). Band intensities were quantified using ImageJ (60).

### Differential scanning fluorimetry

Differential scanning fluorimetry (DSF) was performed as described previously (34). Virion and control (lysozyme) samples were resuspended in one of three buffers: Nanopure sterile water (pH 7.1), virus suspension buffer (50 mM sodium acetate, 8 mM magnesium acetate, pH 4.5), or 100 mM phosphate buffer (pH 7.2). Each experiment was conducted in triplicate, three independent times. DSF data were analyzed and plotted as described previously (34, 35).

### MALDI-TOF

Matrix-assisted laser desorption ionization–time of flight (MALDI-TOF) MS was performed as described previously (35). Purified virion preparations were diluted to 1 mg/ml in 25 mM Tris-HCl, 1 mM EDTA buffer. A 30 μl aliquot (∼30 μg of virions) was digested with trypsin (Pierce, Thermo Fisher) at 1:100 [wt/wt] for varying times at 25°C (61). Samples were analyzed on an AB Sciex TOF/TOF 5800 mass spectrometer with α-cyano-4-hydroxycinnamic acid matrix. Data were processed using AB Sciex Data Explorer software with standard parameters. Peptide fragments were identified using the UCSF Protein Prospector MS-Digest function.

### Progeny analysis

Analysis of viral progeny derivatives was performed as described previously (62). To examine the expression of CP in agroinfiltrated (source) versus non-infiltrated (sink) cells, agrocultures containing either WT BMV or pB1 + pB2 + pB3/P10 were spot-infiltrated into *N. benthamiana* leaves (Fig. 8A). Leaf discs (∼4 mm) were collected from source and sink regions (∼20 mm apart) at 4, 7, and 10 dpi and ground in liquid nitrogen. Total protein was extracted with Pierce Plant Total Protein Extraction Kit (Thermo Fisher Scientific, A44056) and was analyzed by SDS-PAGE and Western blotting. Anti-BMV CP was used at a ratio of 1:1000 (63), and goat anti-rabbit IgG-HRP (1706515) was purchased from Bio-Rad Laboratories and used at a ratio of 1:3,000. For the RT-PCR analysis, total RNA was extracted from specified areas of the leaves using a Qiagen RNeasy Micro kit (Qiagen, 74004). cDNA was synthesized from 100 ng RNA using iScript cDNA Synthesis Kit (BioRad, 1708890). PCR was performed using 2.5 μL of cDNA template and NEB Phusion Hot Start Flex DNA polymerase according to the manufacturer’s instructions, with the following BMV-specific primers: BMV RNA1, forward 5′-CAGTGAGAGAGGTAGAGGAGATAG-3′ and reverse 5′-GTCCCATAGGTGACGAAGATTAG-3′; BMV RNA2, forward 5′-GGTTCAACAGTTCACCGATAGA-3′ and reverse 5′-CTGAGCACTCCCGATATTCATT-3′; BMV movement protein, forward 5′-CTCGTTCGTACCACAGATAGC-3′ and reverse 5′-CGAGAAGATACCCATGATCCAC-3′; and BMV capsid protein (CP), forward 5′-GCTGCCGCTCGTAGAAAT-3′ and reverse 5′-CAGCAACGCTAGGAAGAAGT-3′.

## Authorship contribution statement

**Antara Chakravarty:** Conceptualization, Methodology, Investigation, Visualization, Data Curation, Writing- Original Draft, Writing-Review & Editing. **A. L. N. Rao:** Conceptualization, Supervision, Funding acquisition, Writing-Review & Editing.

## Acknowledgements

We thank Matthew Dickson at the Center for Advanced Microscopy and Microanalysis facility at UC Riverside (UCR) for assisting with imaging the virions; Jie Zhou at the Analytical Chemistry Instrumentation Facility at UCR for helping with MALDI-TOF analysis; and the Institute for Integrative Genome Biology at UCR for allowing us to use the facility for the DSF experiment. Schematics in Figs. 2, 8a, and 9 were created in BioRender and are published here under a CC-BY-NC-ND license. Open access to this article does not include the use of images created in BioRender.

## Data availability

All data supporting the findings of this study are included in the figures.

